# Kv11.1 (hERG) Protein Interaction Networks Connect Endocytic Trafficking to Polygenic Influences on Cardiac Repolarization

**DOI:** 10.64898/2026.01.27.701872

**Authors:** Christian L. Egly, Lea Barny, Suah Woo, Devyn Mitchell, Matthew Ku, Björn C. Knollmann, Lars Plate, Brett Kroncke

## Abstract

Polygenic scores (PGS) summarize the combined effects of common single-nucleotide polymorphisms and contribute to predictions of disease severity, but biological consequences linked to these common variants remain poorly defined. Here, we focused on polygenic liability for a measurable electrophysiologic trait (the QT interval). Prolonged QT interval, measured on patient electrocardiograms, is associated with an increased risk for cardiac arrhythmia. We investigated human induced pluripotent stem cell cardiomyocytes (hiPSC-CMs) from donors with extreme PGS (i.e., high and low) related to QT interval duration. We paired global proteomics with multiplexed affinity purification mass spectrometry (AP-MS) centered on Kv11.1 (hERG), a major determinant of QT-interval repolarization. Global proteomics indicated increased mitochondrial protein abundance in high-PGS cardiomyocytes, but this did not explain the Kv11.1 interactome. In high-PGS cells, Kv11.1 showed increased associations with myosin motor proteins and endosomal recycling machinery, consistent with altered (and potentially increased) recycling/trafficking dynamics rather than trafficking deficiency observed with most pathogenic Kv11.1 variants. This proof-of-concept study underscores a framework for linking polygenic factors to tractable biological consequences by combining patient-specific hiPSCs, proteomics and affinity-purification. Linking polygenic scores to changes in protein networks provides testable mechanisms that can be applied across many diseases.

**Significance Statement:** Polygenic scores (PGS) predict disease risk, but how biological pathways are influenced by these common variants remains difficult to define. We generated human induced pluripotent stem cells from individuals with extreme high- and low-PGS for QT interval, a key electrocardiographic measure linked to arrhythmia risk. By combining global proteomics and interactomics for a common ion channel involved in regulating the QT interval (Kv11.1) we found mechanisms that are influenced by common genetic traits in patients. Our work connects polygenic scores to pathway-level molecular mechanisms in human cells and provides a general framework for uncovering how complex genetic architecture drives disease-relevant biology.

**Classification:** Biological Sciences; Medical Sciences

## Introduction

Long QT Syndrome (LQTS) is an inherited cardiac disorder characterized by a prolonged ventricular repolarization (elongated QT interval on the electrocardiogram), which increases the risk of fatal arrhythmias. While 90% of genotyped LQTS cases are attributed to rare, high-impact mutations in the cardiac ion channel genes *KCNQ1*, *KCNH2*, and *SCN5A*,^1–3^ genome-wide association studies (GWAS) identify many additional genetic loci which explain ∼8-10% of QT interval variability and susceptibility to the drug induced form of the disease (diLQTS).^4–8^ Yet translating these statistical associations into biology is challenging: most QT-associated GWAS loci fall in noncoding regions with subtle, distal regulatory effects.^9–11^ To address this, we developed a system to interrogate the proteomic landscape of human induced pluripotent stem cell–derived cardiomyocytes (hiPSC-CMs) from donors with extreme QT interval polygenic scores. We hypothesize that polygenic liability is not diffuse, but rather converges on specific, actionable biological pathways that control repolarization.

Heterologous expression systems and animal models typically isolate single genetic factors in a uniform genetic background, which fails to recreate the complex genetics of human patients. Additionally, these models have several limitations including either lacking a cardiac context *in vitro* or species differences *in vivo*.^12^ In light of these limitations, human induced pluripotent stem cells (hiPSCs) have emerged as a useful tool to model patient-specific diseases.^13–15^ Studies demonstrate hiPSC-derived cardiomyocytes (hiPSC-CMs) recapitulate human disease with electrophysiological phenotypes that mirror patient QT interval prolongation.^16–19^

Previously, we generated hiPSC-CMs from patients with QT interval polygenic scores (QT-PGS) at the extreme ends of the distribution (above the 99^th^ percentile and below the 1^st^ percentile) and a reference line representing the mid-range.^20,21^ The sensitivity of these cardiomyocytes to pharmacological block of the Kv11.1 potassium channel mirrored the QT-PGS: high-QT-PGS cells were more sensitive to blockade.^21,22^ These differences point to *KCNH2*, which encodes the pore-forming subunit of the Kv11.1 channel or *human Ether-a-go-go-related gene* (hERG). Yet, biological mechanisms linking QT-PGS and QT duration remain unknown. Given that *KCNH2* is central to both congenital^23^ and drug-induced LQTS,^24,25^ we hypothesized that the cumulative burden of background genetic variation exerts its effect by reshaping the macromolecular environment of the Kv11.1 channel. Here, we utilized affinity purification-mass spectrometry (AP-MS) to map the dynamic Kv11.1 protein interaction network (interactome) across polygenic extremes.^26–29^ By profiling these interactions in a human cardiomyocyte context, we identify novel protein partners and pathways that translate polygenic risk into functional susceptibility.

## Results

### Investigating molecular mechanisms impacting QT interval using proteomics and interactomics in hiPSC-CMs

Our primary objective was to use hiPSC-CMs derived from patients with different genetic backgrounds to identify pathways responsible for QT interval variability in patients. We generated hiPSCs from individuals at the extremes of the QT-PGS distribution, plus a reference line (**Fig. 1A**).^20^ Given our prior observation that E-4031 sensitivity tracks with QT-PGS, we analyzed global proteome variation (**Fig. 1B**) and Kv11.1 (*KCNH2*) interaction networks using AP-MS (**Fig. 1C**).^21^ This strategy could uncover altered channel organization, localization, trafficking, and signaling.

**Figure 1.**
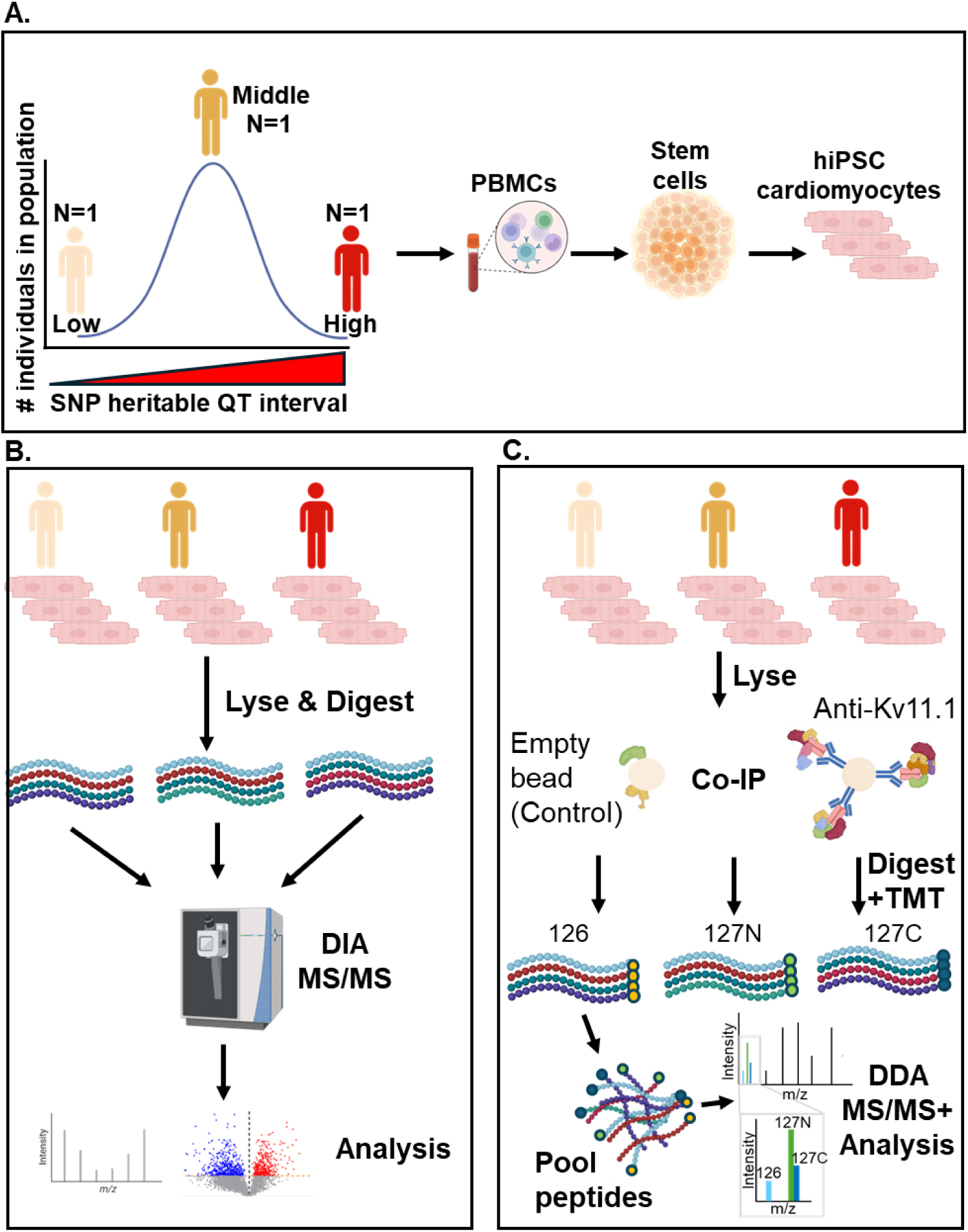
Overview of study design, sample processing, and multi-omics profiling workflow. **(A) Cell line generation**: hiPSCs were derived from patients at upper or lower extremes of QT-PGS, plus one mid-range reference donor, and differentiated into hiPSC-CMs for proteomic analysis. **(B) Global proteomics**: hiPSC-CMs were lysed, digested and analyzed by label-free data-independent acquisition (DIA) mass spectrometry. **(C) Affinity purification/coimmunoprecipitation coupled with mass spectrometry**: Kv11.1 complexes were enriched by co-immunoprecipitation using an anti-Kv11.1 antibody or bead-only controls, followed by digestion and analysis by mass spectrometry using data-dependent acquisition (DDA). Peptides were pooled within run as indicated and analyzed to characterize Kv11.1 interactors and their quantitative differences across QT-PGS groups.

### Global proteomics identifies QT-PGS-associated expression differences in hiPSC-CMs

We used previously described differentiation and maturation protocols^30–32^ to generate >95% pure, spontaneously contracting ventricular hiPSC-CM monolayers. To quantify protein abundances from low-, middle-, and high-QT-PGS cells, we performed data-independent acquisition mass spectrometry. This global proteomic analysis identified 6,430 proteins across all samples (**Supplemental Figure S1A**). Of these proteins, 3,838 (61%) were observed in all three lines (low, middle, and high QT-PGS) at least three times (**Supplemental Figure S1B**). For differential analysis, we filtered proteins by at least three observations in each QT-PGS group and two independent differentiations, yielding 3,666 comparable proteins (**Supplemental Figure S1C&S1D**). Using principal component analysis (PCA) to group samples, we found PC1 primarily reflected batch effects due to differentiation. Comparatively, PCA using PC2 and PC3 consistently separated samples by QT-PGS group, with no apparent outliers (**Figure 2A, Supplemental Figure S1E**). Collectively, these observations mirror previous reports that donor identity drives most expression level variability.^33,34^

**Figure 2.**
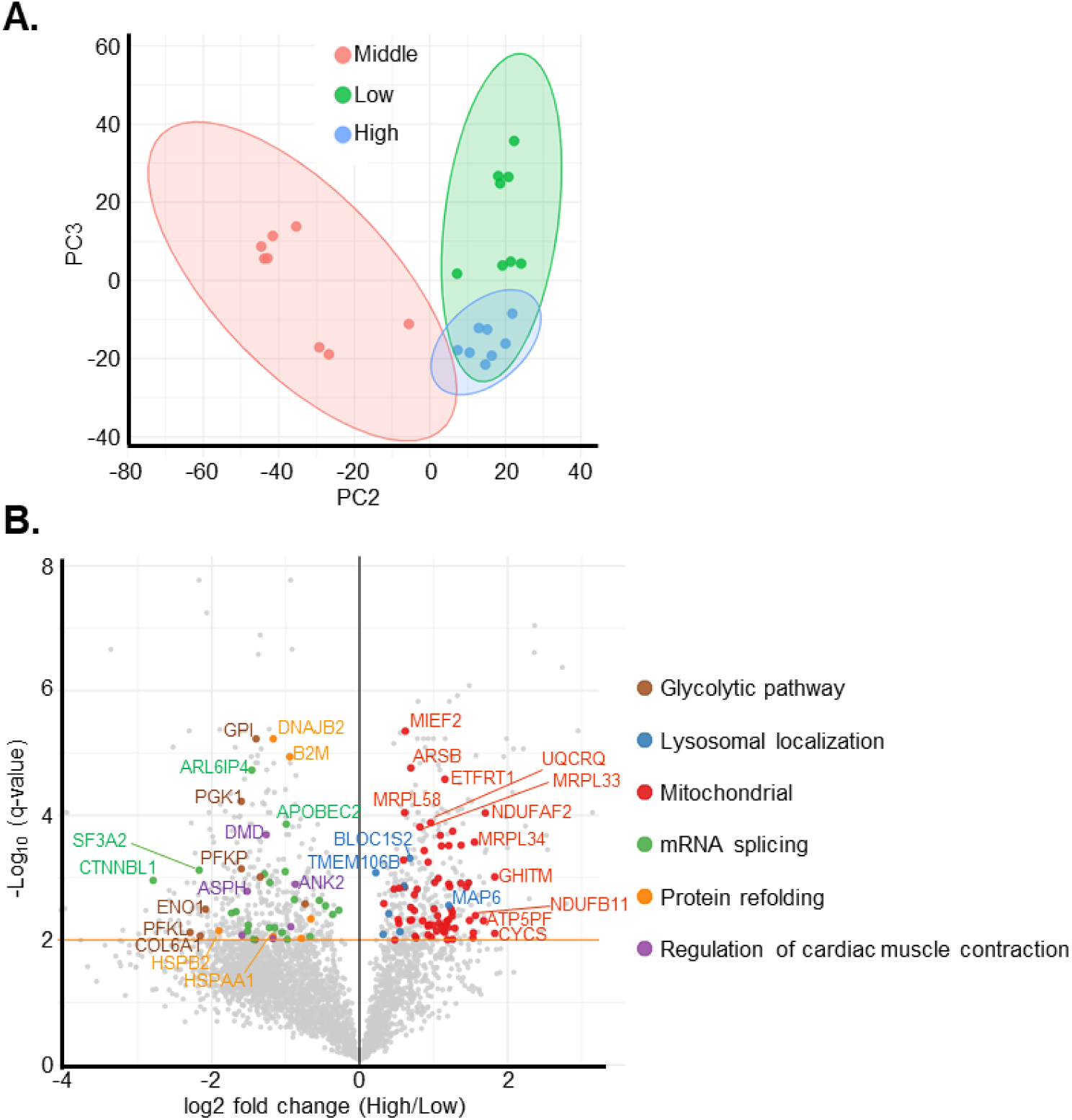
Global proteomics identifies QT-PGS-associated proteome differences in hiPSC-cardiomyocytes. **(A)** Principal component analysis plot showing principal components 2 and 3 for the global proteomics after filtering for proteins identified in at least three samples and two separate differentiations. **(B)** Volcano plot showing log_2_ fold change versus -log_10_ adjusted P values for low- and high-QT-PGS groups. The orange dashed line indicates the FDR threshold for significance (FDR < 0.01; two-stage step-up procedure of Benjamini, Krieger, and Yekutieli). Red = upregulated, blue = downregulated, black = not significant. Samples = 8 lysates per QT-PGS group from at least 2 separate differentiations (Low = 2 differentiations, High = 3 differentiations).

Our global protein analysis revealed 742 proteins with altered expression between high and low QT-PGS (**Figure 2B**), 1,838 between high and middle, and 623 between middle and low (**Supplemental Figure S2A-C**). The middle-QT-PGS line showed a higher proportion of upregulated proteins relative to the other two lines (**Supplemental Figure S1F**). We used the Database for Annotation Visualization and Identification (DAVID) to highlight pathways that differed between cell lines. High QT-PGS cells displayed elevated mitochondrial protein expression: 13 of the top 20 enriched GO terms related to mitochondrial function, translation, or transport, and included iron-sulfur cluster assembly and iron homeostasis (**Figure 3A-B**). These features could indicate an elevated metabolic demand in high-QT-PGS hiPSC-CMs.^35^ In comparison, protein refolding (*HSP90AA1*, *DNAJA1*) was enriched in low- and middle-QT-PGS cells compared to high and suggest a reduced protein folding capacity in the high-QT-PGS group (**Figure 3A-B, Supplemental Figure S2D-E**). Ribosomal translation and mRNA handling pathways were also elevated in the low- and middle-QT-PGS groups (**Supplemental Figure S2D-E**). These data suggest variable protein expression reflecting underlying genetic backgrounds associated with QT liability.

**Figure 3.**
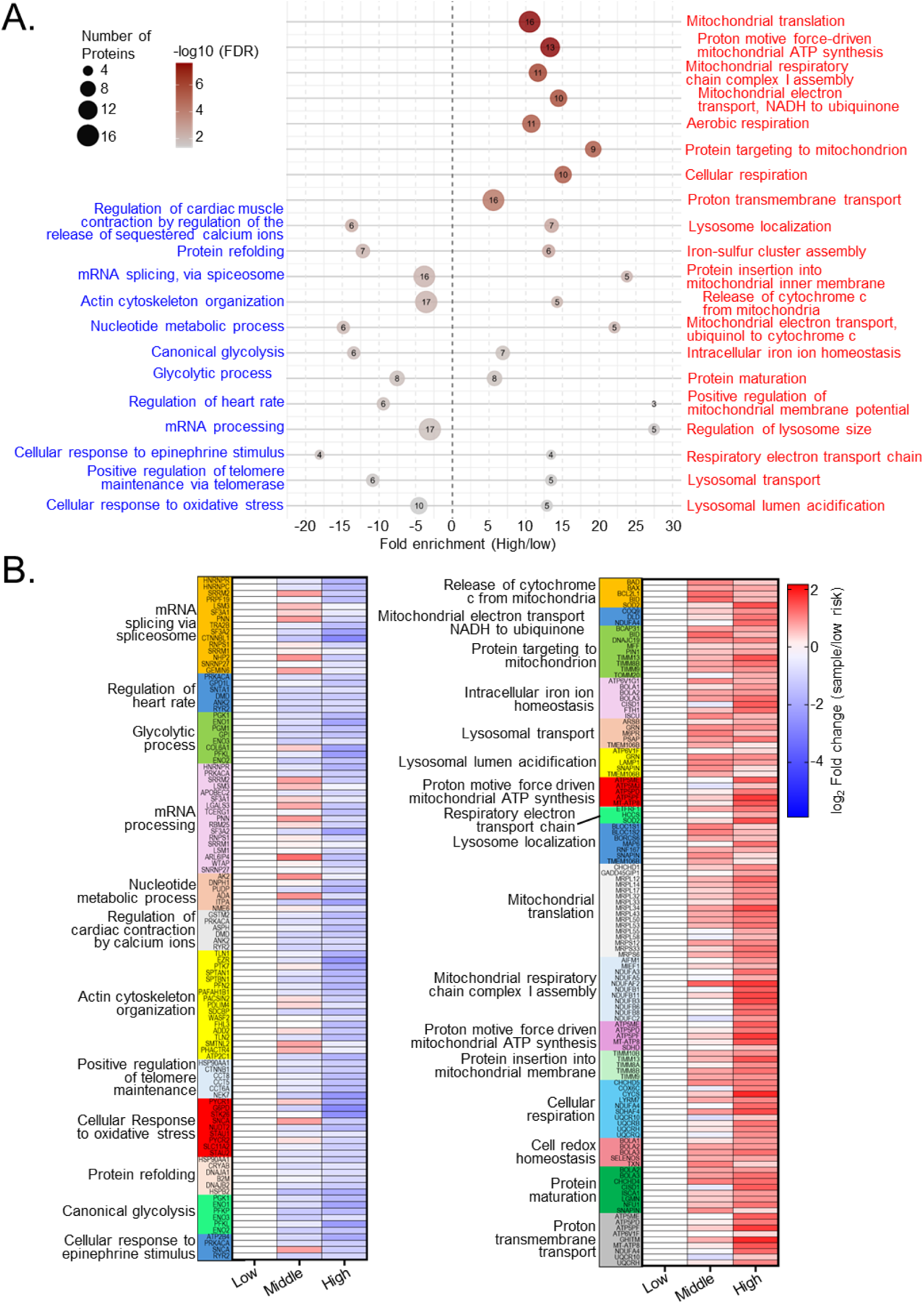
Pathway enrichment and QT-PGS comparisons of protein expression in hiPSC-CMs. **(A)** Bubble plot of DAVID-enriched gene ontology (GO) terms, highlighting pathways increased in high- (red; right of dotted line) or low-QT-PGS (blue; left of dotted line) groups. Numbers within bubbles indicate quantity of enriched proteins matching each GO term. **(B)** Heatmaps showing log_2_ fold change in protein abundance (normalized to low-QT-PGS) for proteins assigned to each GO term. Rows represent proteins; columns represent samples or group means (as indicated). Overlapping genes were reduced to minimize redundancy. N = 8 protein samples per group. Significance was assessed at FDR < 0.01 using a two-stage step-up procedure (Benjamini, Krieger, and Yekutieli).

### AP-MS reveals QT-PGS-dependent remodeling of the Kv11.1 interactome

Given the strong association between QT-PGS and drug-induced LQTS risk, and our prior observation that field potential duration correlates with QT-PGS in these cells, we hypothesized that Kv11.1 regulation or interactions might differ across genetic backgrounds. We used AP-MS to characterize the Kv11.1 interactome in each cell line. For quantification and interactor abundance comparisons, we used tandem mass-tag labeling to multiplex up to 18 samples in single mass spectrometry runs and performed global median normalization across 3 mass spectrometry runs (**Supplemental Figure 3A-B,** n = 6-8 biological replicates per condition). Kv11.1 was significantly enriched compared to bead-only control for all QT-PGS cell lines (**Supplemental Figure 3C**). We identified 319 Kv11.1 interactors significantly enriched over bead-only controls in at least one cell line (**Figure 4A-B**). We manually assigned GO terms to these interactors using Uniprot. Notably, the low and middle QT-PGS lines showed ∼54% stronger Kv11.1 interactions with mitochondrial proteins (P = 0.002), a sharp contrast to the higher mitochondrial protein abundance in high-QT-PGS lysates (**Figure 4C**). This disparity indicates that these localization-dependent Kv11.1 interactions are not accounted for by protein expression differences. Several mitochondrial protein interactors (NDUFA10, ATP51B, COX4I1, COX6C) overlap with our prior findings of Kv11.1 interactions in HEK cells.^36^ Complete heatmaps for the assigned GO terms are provided (**Supplemental Figure 4)**. We identified 43 interactors as significantly enriched across all three cell lines, defining a core Kv11.1 interactome (**Supplemental Figure 5A-D**).

**Figure 4.**
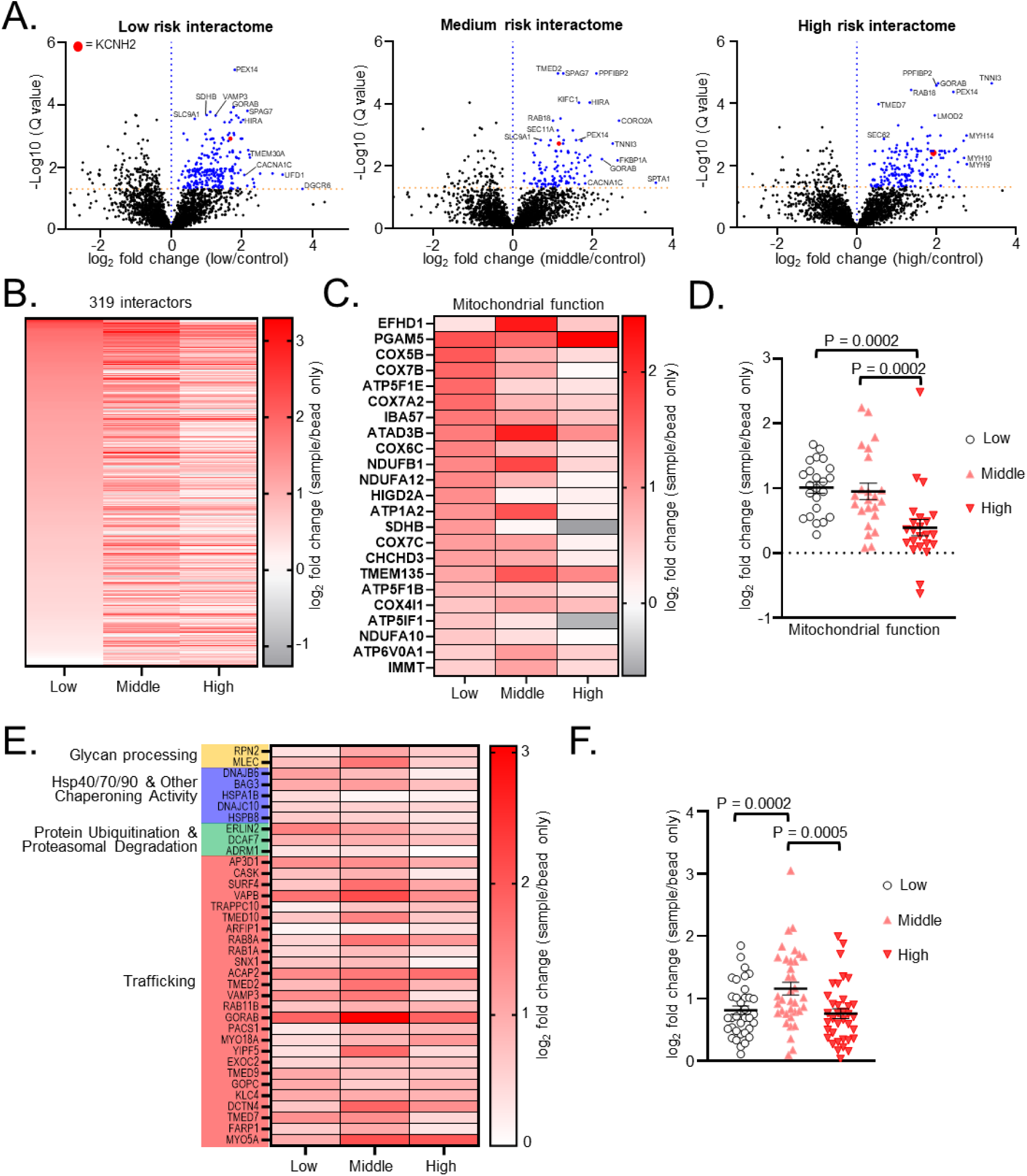
Differential protein abundance of Kv11.1 interactors across QT-PGS groups. **(A)** Volcano plots showing log_2_ fold change versus -log_10_ adjusted P value for each QT-PGS group compared to bead-only background controls. Orange dashed line indicates FDR < 0.05 (two-stage step-up with Benjamini, Krieger, and Yekutieli, n = 6-8 biological replicates per condition). Blue dots are statistically enriched Kv11.1 interactors relative to bead-only background controls. Red highlight is the *KCNH2*/Kv11.1 bait. **(B)** Heatmaps of significantly enriched protein interactors relative to bead-only controls. Rows represent proteins; columns represent samples or group means (as indicated). **(C)** Heatmap of mitochondrial-function proteins (GO terms manually assigned using UniProt) showing relative abundance across groups. **(D)** Dot plot of log_2_ fold change (sample/bead-only) for mitochondrial proteins across groups. **(E)** Heatmap of proteins associated with quality control and trafficking. **(F)** Dot plot of log_2_ fold change comparing all ER quality control and trafficking across groups. Statistics for dot plots were performed using one-way ANOVA with post hoc Tukey’s multiple comparisons test. Data shown as the mean ± SEM.

Kv11.1-protein interactions with glycan-processing machinery, heat shock proteins (HSP40/70/90), the ubiquitin-proteasome system, and general trafficking factors were unchanged between high- and low-QT-PGS groups. This argues against a major trafficking defect in high-QT-PGS cells (**Figure 4E&F**). In contrast, the middle-QT-PGS line showed 37% higher interactions with trafficking-related proteins than either the low or high-QT-PGS, suggesting a mild trafficking deficiency. However, the quantity of Kv11.1 protein was not altered in hiPSC-CM lines, which contrasts with trafficking-defective Kv11.1 variants that show large reductions in Kv11.1 protein abundance compared to wild-type Kv11.1 (**Supplemental Figure 3C**).^36^

From 319 Kv11.1 interactors, DAVID enrichment analysis highlighted cardiomyocyte-relevant processes, including cytoskeleton organization, sarcomere organization, and muscle contraction (**Figure 5A**, **Supplemental Figure 6**). Plasma-membrane Kv11.1 interactions – those that support membrane localization, sodium export (concomitant with potassium influx) and homeostasis, and repolarization – were lower in high-QT-PGS, consistent with decreased surface retention of Kv11.1 (**Figure 5B**). DAVID analysis showed increased enrichment for mitochondrial protein interactions (cellular respiration, proton transmembrane transport, and mitochondrial electron transport cytochrome c to oxygen) in the low-QT-PGS group, which matched our manual annotations (**Supplemental Figure 6**).

**Figure 5.**
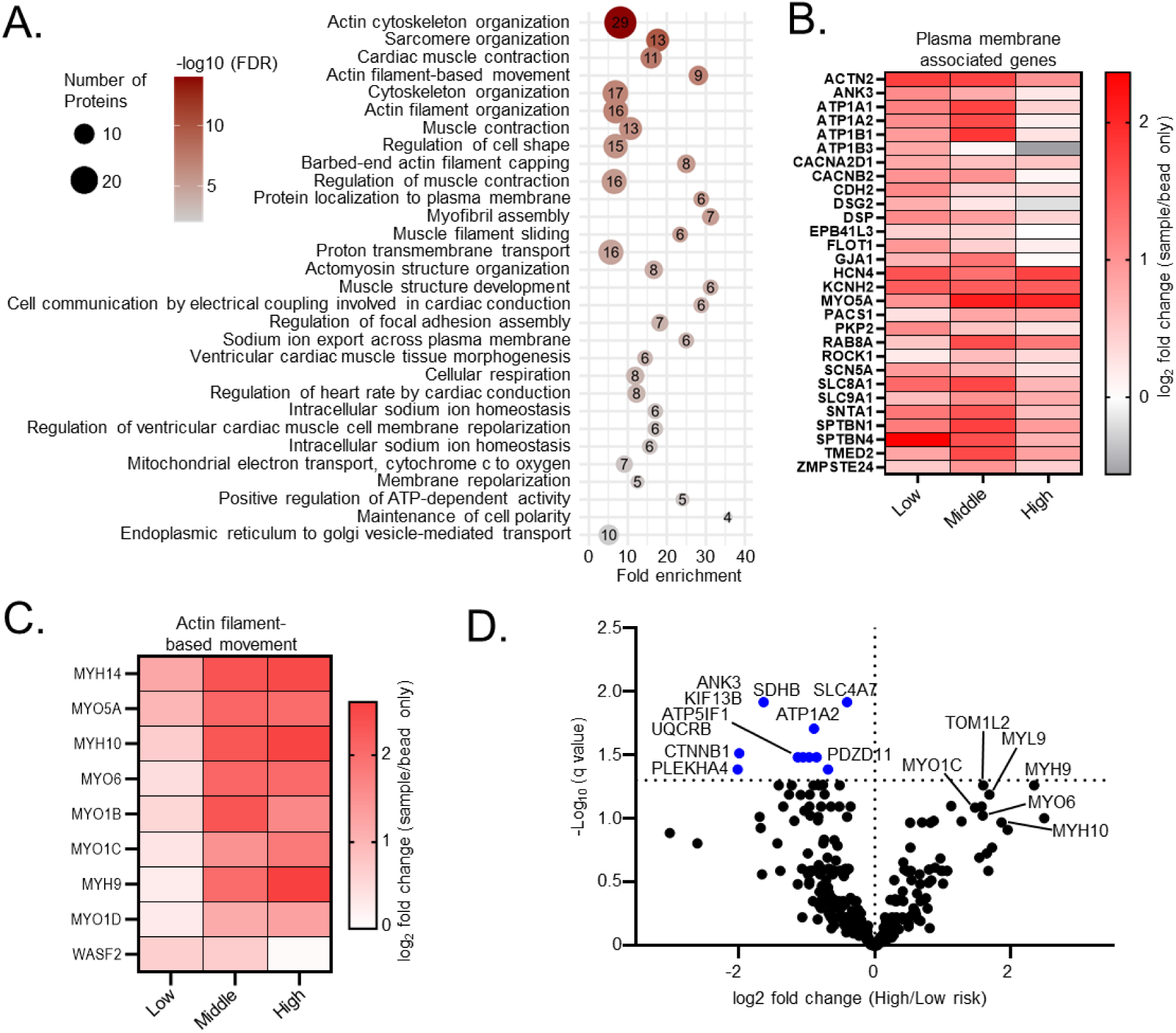
DAVID enriched GO terms of Kv11.1 interactors. **(A)** Bubble plot of the top 30 GO terms (ranked by significance) for all 319 Kv11.1 interactors. **(B)** Heatmap of log2 fold changes for plasma membrane-associated proteins drawn from top GO terms. **(C)** Heatmap of log2 fold changes for actin filament-based movement proteins comparing QT-PGS levels. **(D)** Volcano plot comparing high- vs low-QT-PGS Kv11.1 interactors, showing log_2_ fold changes and -log_10_ adjusted P values.

Kv11.1 association with actin filament-based motility proteins, which includes unconventional myosin motors, was increased in the middle- and high-QT-PGS lines (**Figure 5C**). Compared to low-QT-PGS cells, high-QT-PGS cells had elevated interactions with MYL9, MYH9, MYO6, and MYO1C, and endocytosis related protein TOM1L2 which facilitates MYO6 mediated endocytosis (**Figure 5D**).^37^ These results are consistent with more active, actin-driven endosomal recycling/internalization in high QT-PGS cells.

Individual Kv11.1 protein interactions significantly elevated in low-QT-PGS cells included mitochondrial proteins, cytoskeletal and adhesion proteins, and ion transporters (**Figure 5D**). Enriched mitochondrial interactors included the cytochrome b-c1 complex subunit 7 (*UQCRB*), mitochondrial ATPase inhibitor (*ATP5IF1*) and succinate dehydrogenase iron-sulfur subunit (*SDHB*). Membrane-associated cytoskeletal and adhesion proteins included the pleckstrin homology domain containing family A member 4 (*PLEKHA4*), PDZ domain-containing protein 11 (*PDZD11*), catenin beta-1 (*CTNNB1*), and Ankyrin-3 (*ANK3*). Ion transporters with increased association in low-QT-PGS cells included the sodium/potassium ATPase transporter (*ATP1A2*) and sodium bicarbonate transporter 3 (*SLC4A7*). These interactor shifts showed weak correlation with global protein abundance, further suggesting Kv11.1 interactomes are not driven by expression differences (**Supplemental Figure 7**).

In low-QT-PGS hiPSC-CMs, Kv11.1 preferentially associated with plasma membrane-related proteins, including ion channels, cytoskeletal membrane proteins, and cell adhesion/gap-junction components. This pattern suggests that Kv11.1 from low-QT-PGS cells is more frequently anchored at the plasma membrane. Kv11.1 in high-QT-PGS cells exhibited strengthened associations with actin-based myosin trafficking machinery and Rab-family GTPases that mediate endosomal recycling. These data highlight pathways that may contribute to QT interval control, shifting Kv11.1 channels from membrane-anchored complexes toward endosomal trafficking machinery, summarized in **Figure 6**.

**Figure 6.**
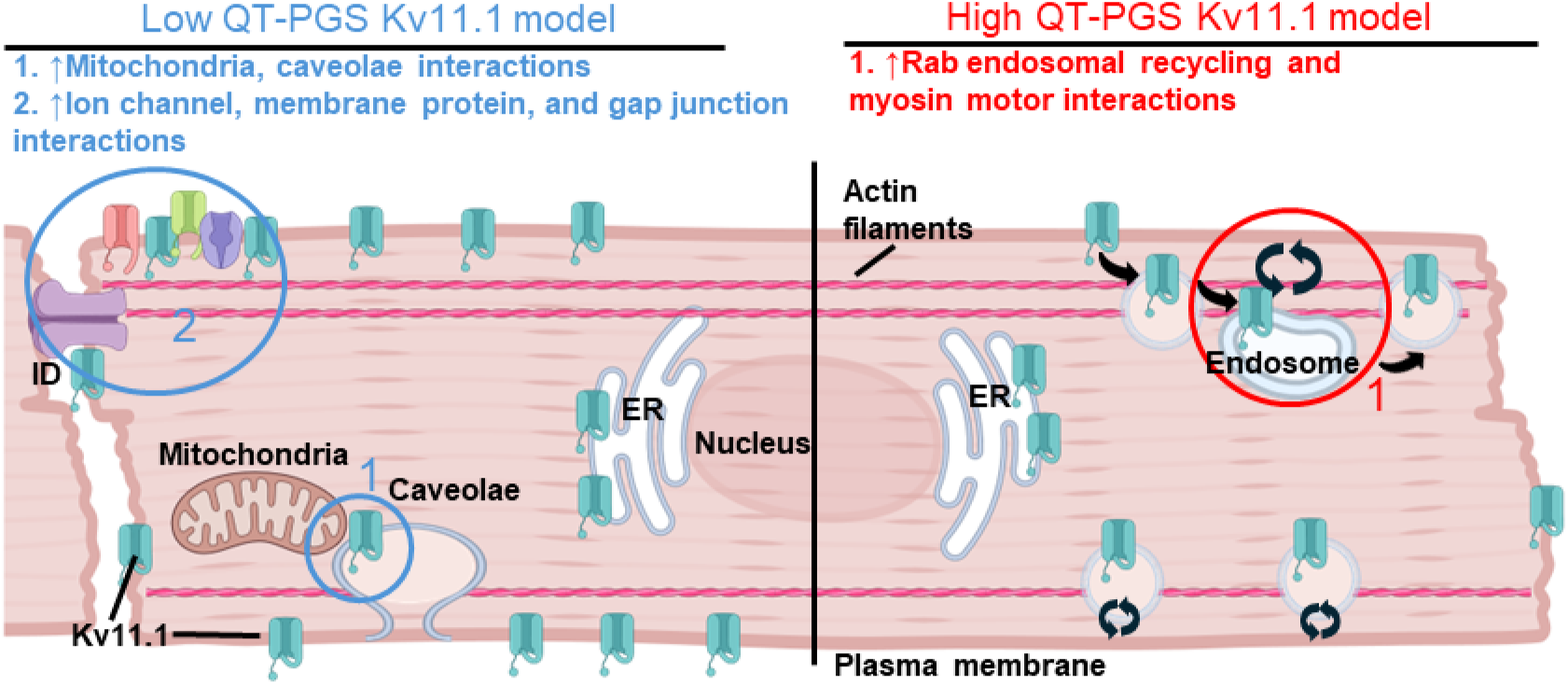
Cellular model of Kv11.1 interactomics and QT-PGS comparisons. Schematic model summarizing Kv11.1 interactions and localization across subcellular compartments and their PGS-associated differences between low and high. ID – intercalated disk, ER – endoplasmic reticulum.

## Discussion

In this study, we investigated how polygenic QT interval influences cardiomyocyte biology by combining global proteomics with Kv11.1 interactome profiling across hiPSC-cardiomyocytes (CM) stratified by QT interval PGS. High QT-PGS hiPSC-CMs displayed a pronounced shift toward mitochondrial protein enrichment with 13 of the 20 top enriched GO terms relating to mitochondrial function, suggesting elevated metabolic demand in these cells. Conversely, low and middle QT-PGS cells showed enrichment for protein refolding machinery as well as ribosomal and mRNA handling pathways, pointing to enhanced proteostatic capacity. Interactome profiling identified 319 Kv11.1-associated proteins, 43 of which formed a core interactome conserved across all lines. Paradoxically, despite their lower overall mitochondrial protein abundance, low and middle QT-PGS cells exhibited approximately 54% stronger Kv11.1-mitochondrial protein interactions, suggesting that these associations are driven by subcellular localization rather than expression levels alone. High QT-PGS cells instead showed reduced plasma membrane-associated Kv11.1 interactions coupled with elevated associations with myosin motors and endocytic machinery. Importantly, core trafficking and folding machinery interactions remained unchanged across lines, and Kv11.1 protein levels themselves were equivalent, arguing against a primary trafficking defect and instead implicating altered Kv11.1 localization or functional microenvironment as the mechanistic link between polygenic burden and arrhythmia susceptibility. These pathway-level shifts may reflect altered repolarization reserve in cells with high polygenic QT liability.

We previously reported 570 Kv11.1 interactors in human embryonic kidney (HEK-293) cells.^36^ Here, we identified 318 Kv11.1 interactions in hiPSC-CMs with 78 (∼25%) proteins overlapping across studies, comparable with other interactome studies.^38^ Enriched GO-terms highlighted actin cytoskeleton organization and actin filament organization as top Kv11.1 interacting categories in HEK and hiPSC-CM cells. However, proteosomal interactions were enriched in HEK-293 cells but minimal in hiPSC-CMs.

Our global proteomics data showed higher mitochondrial protein abundance in high-QT-PGS cells, consistent with previous reports.^26,36^ These proteins include components of complex I assembly (NDUF family), mitochondrial ribosomes (MRPL/MRPS), cellular respiration (COX/UQCR), and mitochondrial membrane formation (TIMM). We also observed increased plasma membrane and caveolar interactions with Kv11.1 in the low-QT-PGS group suggesting these Kv11.1-mitochondrial complexes form near the plasma membrane. Prior reports in cardiac tissue have reported mitochondria form close interactions with caveolae near the cellular surface.^39,40^ Thus, enrichment of caveolar and mitochondrial membrane components in low/middle-QT-PGS cells supports the presence of mitochondrial-associated complexes near the plasma membrane with potential roles in ion channel function. Intriguingly, one study found that increased mitochondrial interactions with caveolae at the plasma membrane were associated with decreased cardiac damage.^40^ This suggests these interactions have a protective role and preserve ion channel function at the cellular membrane when comparing low- to high-QT-PGS.

Changes in endosomal recycling and trafficking shift towards the lysosome rather than the proteasome in high-QT-PGS cardiomyocytes. Kv11.1 interactions in high-QT-PGS cells were elevated for Myosin V (*MYO5A*, antegrade) and Myosin VI, (*MYO6*, retrograde/endocytic), Rab GTPases (Rab18 and Rab8A), and Tom1L2, which can interact with MYO6 to transport vesicle cargo to the lysosome;^37^ MYO6 also mediates endocytosis of the cystic fibrosis transmembrane conductance regulator.^41^ We identified Rab11B, which is a well-known interactor of Kv11.1 responsible for slow endosomal recycling.^42^ This, together with increased Kv11.1 interactions with other myosin motor proteins may suggest increased endosomal recycling from the membrane to intracellular endosomes and the lysosome rather than proteasome.

This proof-of-concept study underscores the feasibility of using hiPSC-derived cardiomyocytes as a platform to connect polygenic risk with mechanistic pathways. In these patient-derived lines, we found endosomal recycling and interactions with unconventional myosin motors were associated with high-QT-PGS. Limitations in our study include small sample size (N = 1 patient each at low- or high-QT-PGS extremes and one reference line). Future validation across multiple independent lines should identify whether these are common features. Additionally, our global proteomics data revealed differentiation-dependent variability, a well-recognized issue in hiPSC work (**Supplemental Figure 1**).^43^ Nonetheless, enrichment for mitochondrial proteins remained consistent when comparing high-QT-PGS lysates to individual low-QT-PGS differentiations. Additionally, the differentiation and maturation protocol we use produces highly pure cardiomyocytes (>95%), and we verified hiPSC-CMs for spontaneous contraction for all experiments.^30–32^ In addition, unbiased clustering through principal component analysis grouped samples by QT-PGS without outliers, supporting the presence of a robust QT-PGS-related signal despite batch effects.

In conclusion, we have used global proteomics and Kv11.1 interactomics to reveal specific pathways that alter cellular electrophysiology and contribute to QT interval variability. We show that hiPSC-CMs from patients with different QT-PGS do not have major changes in Kv11.1 folding or trafficking, in contrast with Kv11.1 variants.^44,45^ Instead, we found altered interactions with endosomal recycling machinery, particularly unconventional myosin proteins and altered mitochondrial interactions. These findings suggest that polygenic risk operates through distinct mechanisms from monogenic disease, potentially by modulating channel localization and functional microenvironment rather than biogenesis. Further investigation of these pathways may reveal novel determinants of repolarization reserve and identify new therapeutic targets for arrhythmia susceptibility.

## Materials and Methods

### QT polygenic score and iPS Cell Generation

The QT polygenic score (QT-PGS) was derived from two independent sources: one from Nauffal et al. comprising 1,107,630 SNPs, and one from Arking et al. comprising 68 SNPs.^4,6^ The QT-PGS percentile was calculated using the standard deviation from these SNPs from each patient line selected from a previous cohort of 2,899 genotyped participants; We utilized hiPSCs derived from three individuals exhibiting either extreme high (>95% for each QT-PGS) or low (<1% for each QTPGS) QT-PGS and a reference “middle” line (∼50% for each QT-PGS). Low-and high-QT-PGS individuals were non-Latino, white males over 65 without ventricular arrhythmia-related phenotypes. The reference line was a non-Latino, white female over 65. Peripheral blood mononuclear cells were reprogrammed into hiPSCs using non-integrating episomal vectors following established protocols and characterized in our previous study.^20^

### Differentiation into Cardiomyocytes and maturation protocols

HiPSC cardiac differentiation and maturation methods have been previously published.^31,32^ Briefly, hiPSCs were plated on 6-well plates with 1:200 matrigel and cultured in StemMACS™ iPS-Brew XF Basal Medium (Militenyi Biotec, 130-107-086). Cardiac induction was started at 65-70% confluency using a standard monolayer differentiation method involving small molecules CHIR99021 on day 0-2 (6 µmol/L, Selleck Chemicals) and IWR-1 from day 3-5 (5 µmol/L, Sigma Aldrich).

Metabolic selection was performed from days 10-16 using glucose-free RPMI1640 (catalog no. 11879; Life Technologies). On day 15, hiPSC-CMs were dissociated and replated on 1:200 Matrigel-coated plates in glucose-free media supplemented with 10% FBS. From day 16 to day 30, hiPSC-CMs were treated with 100 nmol/L triiodothyronine (Sigma) and 1 µmol/L Dexamethasone (Cayman) every other day in RPMI1640 with 2% B-27 supplement including insulin (Life Technologies) to improve cardiac maturation as previously described.^31,32^ Cells were maintained in culture until day 30 post-differentiation for subsequent experiments. All hiPSC-CM monolayers were visually analyzed for contraction before lysate collection and subsequent experiments. Cells that did not exhibit spontaneous beating were discarded.

## Global proteomics

### Protein Digestion and Sample Preparation for LC-MS/MS

Protein cleanup and digestion was carried out on the Biomek i5 (Beckman Coulter). 20 µg of protein from each sample was digested (8 samples from each low-, middle-, and high-QT-PGS lines. Proteins were first reduced with fresh dithiothreitol (DTT; 5 mM, Sigma) for 30 min at 37 °C with shaking at 1000 rpm and alkylated with fresh iodoacetamide (20 mM, Sigma) for 15 min at room temperature (in darkness). To quench the alkylation, DTT was added a second time (5 mM) with incubation for 15 min at room temperature (RT)/1000 rpm. Subsequently, proteins were digested on 8 µL’s of magnetic single pot solid phase enhanced sample preparation beads (SP3; Cytiva; 50 mg/mL; 1:1 hydrophilic to hydrophobic beads) and cleaned-up as previously described.^46^ Protein was bound to SP3 beads with 100% ethanol and washed three times with 80% ethanol for clean-up. Proteins were then digested with Trypsin/Lys-C (Thermo Fisher; 1:40 protease to protein ratio) in 50 µL ammonium bicarbonate (ABC; 100 mM; pH 8) at 37 °C / 700 rpm. Digested peptides were removed from SP3 beads after digestion on the Biomek i5 and dispensed manually into low-bind tubes. Formic acid was added to each sample to a final concentration of 2% (v/v) and samples were subsequently dried *in vacuo* using a SpeedVac (ThermoFisher Scientific, SPD111V). Samples were stored dried at -80 °C until use. Peptides were resuspended to a final concentration of 200 ng/µL in buffer A (4.9% ACN, 95% H_2_0, 0.1% FA (v/v/v). Samples were then spun down at 21,000 g for 15 min and then transferred to a fresh low bind eppendorf tube prior to LC-MS/MS analysis.

### LC-MS/MS Analysis

LC-MS/MS analysis was performed using an Exploris480 mass spectrometer (Thermo Fisher) equipped with a Dionex Ultimate 3000 RSLCnano system (Thermo Fisher). Peptides were separated using a 21.5 cm fused silica microcapillary column (ID 100 μm) ending with a laser-pulled tip filled with Aqua C18, 3 μm, 100 Å resin (Phenomenex # 04A-4311). Electrospray ionization was performed directly from the analytical column by applying a voltage of 2.2 kV (positive ionization mode) with an MS inlet capillary temperature of 275 °C and a radiofrequency (RF) Lens of 40%.

A total of 600 ng of peptides per sample were loaded onto a commercial trap column (C18, 5 μm, 0.3x5mm; ThermoFisher Scientific, 160454) using an autosampler. Peptides were then eluted and separated on a 2 h gradient with a constant flow rate of 500 nL/min: 2% B (acetonitrile with 0.1% formic acid, 5 min hold) was ramped to 35% B over 90 min and stepped to 80% over 5 min and held at 80% B for 5 min, followed by a 13 min hold at 4% B to re-equilibrate the column. Between injections, a 45 min column-wash was performed using the following gradient: 2% B (6 min hold) stepped to 5% over 2 min and subsequently ramped to a mobile phase concentration of 35% B over 7 min, ramped to 65% B over 5 min, held at 85% B for 8 min, then returned to 3% B for the remainder of the analysis.

Our data independent acquisition mass spectrometry (DIA-MS) method consisted of 62 MS/MS scan events with 10 *m/z* precursor isolation windows and 1 *m/z* overlaps (30,000 resolution, 1e7 normalized ACG target, 55 ms maxIT, 27% Normalized Collision Energy (NCE), 40% RF lens) taken across a 380-1000 *m/z* mass range. A precursor spectrum (380-1000 *m/z*, 3e6 normalized ACG target, 40% RF lens) was taken at 120,000 resolution with a maxIT of 240 ms. Loop control was utilized with n= 20 to intersperse MS^1^ scans between MS/MS acquisitions. The 66 spectra collected per cycle resulted in a duty cycle time of approximately 4.4 sec.

### Peptide Identification and Quantification

All DIA spectra were analyzed using DIA-NN (version 1.8.1) with a library free search workflow. A spectral library was generated in DIA-NN using the UniProt SwissProt canonical human FASTA database (downloaded on May 1, 2024; containing 20,361 entries) and a contaminant FASTA file (containing 379 entries). Peptides ranging from 7 to 30 amino acids in length were considered, allowing for one missed trypsin cleavage and a maximum of one variable modification (Met oxidation or N-terminal acetylation). N-terminal M excision was enabled and cysteine carbamidomethylation was included as a fixed modification. Precursors between *m/z* 380-1000 with charge states of 1-4 were considered. Additionally, fragment ions between *m/z* 200 and 1800 were included. We then used a single-pass neural network classifier to build an in-silico spectral library from the DIA data and searched the raw files against this library. Smart profiling was used for library generation. Unrelated runs (mass accuracy and scan windows determined separately for different runs), match between runs (MBR), heuristic protein inference, and no shared spectra were selected in the algorithm section of the DIA-NN GUI. Protein inference was made on genes and robust LC (high precision) was used as the quantification strategy. Protein assembly was performed in DIA-NN based on the parsimony principle.

### Data Analysis (Global Proteomics)

A custom R script was developed to process DIA-NN output reports to perform statistical filtering, normalization, and data visualization. As part of this workflow, the diann.dia.qc() function was used to retain peptides with a peptide *q*-value and global protein *q*-value score below 1%. To increase confidence in protein identifications, only proteins supported by at least two peptides were retained for downstream analysis. Contaminant proteins were removed prior to subsequent processing.^47^

Following peptide-level filtering, data were globally median normalized using the medNorm() function. For each sample *i*, a normalization factor F*_i_* was computed as the ratio between the global median of all precursor intensities (M_G) and the median intensity within a particular sample (M_I). The normalized abundance (A ^norm^) for each precursor *j* in sample *i* (A_ij_) was then calculated in the following two steps: (1) F*_i_* = M_G /M_I & (2) A ^norm^ = F*_i_* × A*_ij_*. Protein-level quantifications were then inferred from the filtered and normalized peptides using the max_LFQ() function from the DIA-NN R package^48^.

## Interactome Proteomics

### Covalent Antibody-bead crosslinking

This procedure was adapted from Pankow, et al.^29^ Pierce Protein A/G Magnetic Beads (ThermoFisher) containing a recombinant Protein A/G (∼50.5 kDa) that combines the IgG binding domain of both Protein A and Protein G were used to cross-link anti-Kv11.1 antibodies (Cell Signaling). Stock beads with a concentration of 10 mg/ml were washed four times with 1 mL of Dulbecco’s Phosphate Buffered Saline (DPBS, ThermoFisher). Excess DPBS was removed, followed by the addition of an appropriate amount of anti-Kv11.1 antibodies to the beads to make the final concentration antibody/bead of 60 µg/100 µL, which is equivalent to 42.8 µL of antibody per 100 µL of stock beads. Then, the antibody and beads were gently mixed for two hours at room temperature (20-25°C) on a rotator to allow the binding of the antibody to the beads. Next, the beads were pulled with a magnetic stand and washed twice with 1 mL of freshly made 100 mM sodium borate (pH 9.0, Sigma-Aldrich). To start the cross-linking reaction, beads were resuspended in 1 mL of 100 mM sodium borate (pH 9.0) followed by the addition of dimethylpimelimidate powder (DMP, ThermoFisher) to a final DMP concentration of 25 mM. The bead-antibody mixture was rotated for 30 minutes at room temperature on a rotator, then the beads were pulled with a magnetic stand and the supernatant was discarded. After that, beads were washed once with 1 mL of 200 mM freshly prepared ethanolamine (pH 8.0) (Sigma-Aldrich), resuspended in 1 mL of 200 mM ethanolamine, and incubated for 2 hours at room temperature with gentle mixing. Finally, beads were washed five times with 1 mL of DPBS to remove excess ethanolamine and stored in DPBS with 0.1% Tween-20 (v/v) (Sigma-Aldrich) and 0.02% sodium azide (wt/vol) (Sigma-Aldrich).

### Co-immunoprecipitation (affinity-purification)

HiPSC-CMs from day 30-42 were lysed and protein was harvested for co-immunoprecipitations. The cells were washed with Dulbecco’s phosphate-buffered saline (DPBS, Gibco) without calcium and magnesium. DPBS was removed from 6-well plates and 100 µL/well of ice-cold lysis buffer (same as for global proteomics) was added containing 50 mM Tris pH 7.5, 250 mM NaCl, 1 mM EDTA, 0.5% Igepal-CA-630 (TNI buffer) and 1x protease inhibitor cocktail (Sigma). The flasks were agitated on an orbital shaker for 30 minutes. Cell lysates were scraped from plates into 1.7 mL Eppendorf tubes and sonicated with probe sonication (Qsonica) at 10 watts for two sequences of 2-second-bursts, separated by 7-second pauses while samples were kept on ice. After sonication, samples were centrifuged at 18,000 × g for 30 minutes at 4 °C. Supernatant was collected post-centrifugation and pellets were discarded. Finally, the protein concentration was quantified using a Bicinchoninic acid assay (BCA, Pierce 23225).

Next, 5 mg of lysate representing one biological replicate were incubated for 2 hours at 4 °C with anti-Kv11.1 antibody (Cell Signaling #12889) covalently coupled to protein A/G magnetic beads (Thermo Fisher). The antibody-to-lysate ratio was maintained at 0.1 µg of antibody to 100 µg lysate. Following coimmunoprecipitation in 15 mL centrifuge tubes, complexes were transferred to 1.7 mL Eppendorf tubes. Samples were washed three times (250 µL) with a buffer containing 25 mM Tris-HCl pH 7.5, 150 mM NaCl, and 1 mM EDTA (TN buffer). A magnetic stand was used to separate bead-Kv11.1 protein interactions from the rest of the lysate. After the final wash, all media was removed and antibody-bead/coimmunoprecipitated protein mixture was placed in a –80 °C freezer for 1 hour to facilitate later elution.

For elution, 50 µL of 0.2 M glycine (pH 2.3) buffer was added, and the sample was heated on a heating block (70 °C) for 10 minutes. Sample was collected by separating magnetic beads from eluate and then neutralized by adding 5 µL ammonium bicarbonate (1 M). Samples were stored at –80 °C until proceeding to either western blotting, total membrane staining, or liquid chromatography-tandem mass spectrometry LC-MS/MS analysis.

### Protein digestion and TMT labeling

Eluted samples were precipitated in methanol/chloroform, washed three times with methanol, and air-dried. Protein pellets were then resuspended in 4 μL 1% Rapigest SF Surfactant (Waters) followed by the addition of 10 μL of 50 mM HEPES, pH 8.0, and 32.5 μL of LCMS grade water. Samples were reduced with 5 mM tris(2-carboxyethyl)phosphine (TCEP, Sigma) and alkylated with 10 mM iodoacetamide (Sigma). Next, 0.5 μg of trypsin (Sequencing Grade, Promega, or Pierce) was added and incubated for 16–18 hours at 37 °C with shaking at 700 rpm. Peptide samples were then reacted with tandem mass tag (TMT) 16/18-plex reagents (Thermo Fisher) in 40% vol/vol acetonitrile and incubated for 1 hour at room temperature. Reactions were quenched by adding ammonium bicarbonate (0.4% wt/vol final concentration) and incubated for 1 hour at room temperature. TMT-labeled samples for a given mass spectrometry run were then pooled and acidified with 5% formic acid (Sigma, vol/vol). Samples were concentrated using a SpeedVac and resuspended in buffer A (95% water, 4.9% acetonitrile, and 0.1% formic acid, v/v/v). Cleaved Rapigest SF surfactant was removed by centrifugation for 30 min at 21,100 × g.

### Multidimensional Protein Identification Technology (MudPIT) LC-MS/MS Analysis

LC-MS/MS data-dependent analysis (DDA) for 18-plex TMT samples was performed using an Exploris480 mass spectrometer (Thermo Fisher) equipped with a Dionex Ultimate 3000 RSLCnano system (Thermo Fisher). Peptide samples were loaded using a high-pressure chamber onto a triphasic MudPIT column (prepared as previously described).^49^ Briefly, the MudPIT column was packed with 1.5 cm of Aqua 5 µm C18 resin (Phenomenex # 04A-4299), 1.5 cm Luna 5 µm strong-cation exchange (SCX) resin (Phenomenex # 04A-4398), and 1.5 cm 5 µm C18 resin. Before analysis, the peptide loaded MudPIT column was desalted with buffer A (95% water, 4.9% acetonitrile, 0.1% formic acid v/v/v) for 30 minutes. Peptides were eluted from the first C18 phase onto the SCX resin using a 10 µL injection of buffer A with the following 90-minute gradient: 2% buffer B, composed of 99.9% acetonitrile and 0.1% formic acid v/v (5-minute hold) ramped to a mobile phase concentration of 40% B over 35 minutes, ramped to 80% B over 15 minutes, held at 80% B for 5 minutes, then returned to 2% B for 5 minutes and then held at 2% B for the remainder of the analysis at a constant flow rate 500 nL/min. Peptides were then sequentially eluted from the SCX stationary phase using 10 µL injections of 10, 20, 40, 60, 80, and 100% buffer C (500 mM ammonium acetate in buffer A). The final fraction was eluted with 10% buffer B and 90% buffer C. Fractions were collected using the following 130-minute gradient with a constant flow rate of 500 nL/min: 2% B (6-minute hold) stepped to 5% B over 2 minutes, ramped to 35% B over 92 minutes and stepped to 85% in 6 minutes and held at 85% B for 7 minutes, followed by a 17-minute hold at 2% B. For both gradients, the loading pump was held at 2% B for the duration of the analysis. During buffer pickup and loading for each method, the loading pump was held at a flow rate of 10 µL/minute for 8 mins prior to the valve switch. For 5 minutes before the end of the analytical gradient, the valve was switched to the load position to prepare for sample loading and the flow rate was adjusted from 1 µL/minute to 10 µL/minute for the remainder of the analysis. Peptides were separated using a 20 cm fused silica microcapillary column (ID 100 μm) ending with a laser-pulled tip filled with Aqua C18, 3 μm, 100 Å resin (Phenomenex # 04A-4311). Electrospray ionization was performed directly from the analytical column by applying a voltage of 2.2 kV with an MS inlet capillary temperature of 275 °C and a RF Lens of 40%.

For TMT-DDA acquisition, a 3 second duty cycle was utilized consisting of a full scan (375-1500 *m/z*, 120,000 resolution) and subsequent MS/MS spectra collected in TopSpeed acquisition mode. For MS^1^ scans, the maximum injection time was set to automatic with a normalized AGC target of 300%. Only ions with an MS^1^ intensity above 5e3 with a charge state between 2-7 were selected for MS/MS fragmentation. Additionally, a dynamic exclusion time of 45 seconds was utilized (determined from peptide elution profiles) with a mass tolerance of +/- 10 ppm to maximize peptide identifications. MS/MS spectra were collected with a normalized HCD collision energy of 36, 0.7 m/z isolation window, 150 millisecond maximum injection time, a normalized AGC target of 200%, at a MS^2^ orbitrap resolution of 45,000 with a defined first mass of 110 *m/z* to ensure measurement of TMTPro reporter ions.

### Peptide Identification and Quantification

Peptide identification and TMT-based protein quantification were performed in Proteome Discoverer 2.4 (PD, Thermo Fisher). MS/MS spectra were searched using SEQUEST-HT against a Uniprot human proteome database (released 03/2014 and containing 20,337 entries; supplemented with common MS contaminants and curated to remove redundant proteins/splice isoforms) and a decoy database of reversed peptide sequences. Searches were carried out according to the following parameters: 20 ppm peptide precursor tolerance, 0.02 Da fragment mass tolerance, minimum peptide length of 6 AAs, trypsin cleavage with a maximum of two missed cleavages, dynamic modifications of Met oxidation (+ 15.995 Da), protein N-terminal Met loss (-131.040 Da), protein N-terminal acetylation (+ 42.011 Da), protein N-terminal Met loss and acetylation (+ 89.030), and static modifications of Cys carbamidomethylation (+ 57.021 Da) and TMTPro (N-terminus/Lys, + 304.207 Da). IDs were filtered using Percolator with a peptide level FDR of 1%. The Reporter Ion Quantification node in PD was used to quantify TMT reporter ion intensities. Only reporter ions with an average signal-to-noise ratio (S/N) greater than 10 to 1 (10:1) and a percent co-isolation less than 25 were utilized for quantification. TMT reporter ion intensities were summed for peptides belonging to the same protein, including razor peptides. Proteins were filtered at 1% FDR and protein grouping was performed according to the parsimony principle. Protein identifications and TMT quantification can be found in **Supplemental Table 4**.

### Proteomics analysis

Protein abundances for all channels across each mass spectrometry run were median normalized (N = 3 runs). The median normalized quantifications are included in **Supplemental Figure 3B.** To determine Kv11.1 enriched interactions, data were log_2_ normalized and each protein pulled down with Kv11.1 was compared to the corresponding log_2_ TMT intensity for each protein pulled down in protein A/G beads without antibody. Significance was determined using multiple t-tests with false discovery rate correction by Benjamini, Krieger, and Yekutieli (q < 0.05). For subsequent analysis, we scaled log_2_ protein abundances (sample/beads only) to the Kv11.1 protein within each channel. The individual replicates from separate runs were compiled and averaged to result in the log_2_ fold change for each protein normalized to Kv11.1. We only included proteins identified and quantified in two of three mass spectrometry runs for coimmunoprecipitation results.

## Author Contributions

CLE, LB, LP, and BK contributed to the conceptualization, project administration, visualization, and original draft preparation of the study. CLE and LB were responsible for data curation, formal analysis, methodology, and validation. CLE, BK, and LP contributed to funding acquisition and supervised the study. BK and LP provided resources. CLE, LB and LP provided the software. CLE, SW, LB, DM, and MK conducted the investigation. CLE, LB, LP, BCK and BK contributed to the review and editing of the manuscript.

