## Supplemental figures for "Kv11.1 (hERG) Protein Interaction Networks Connect Endocytic Trafficking to Polygenic Influences on Cardiac Repolarization"

**The Supplementary Materials file includes:**

**Supplemental Figures S1 to S7**

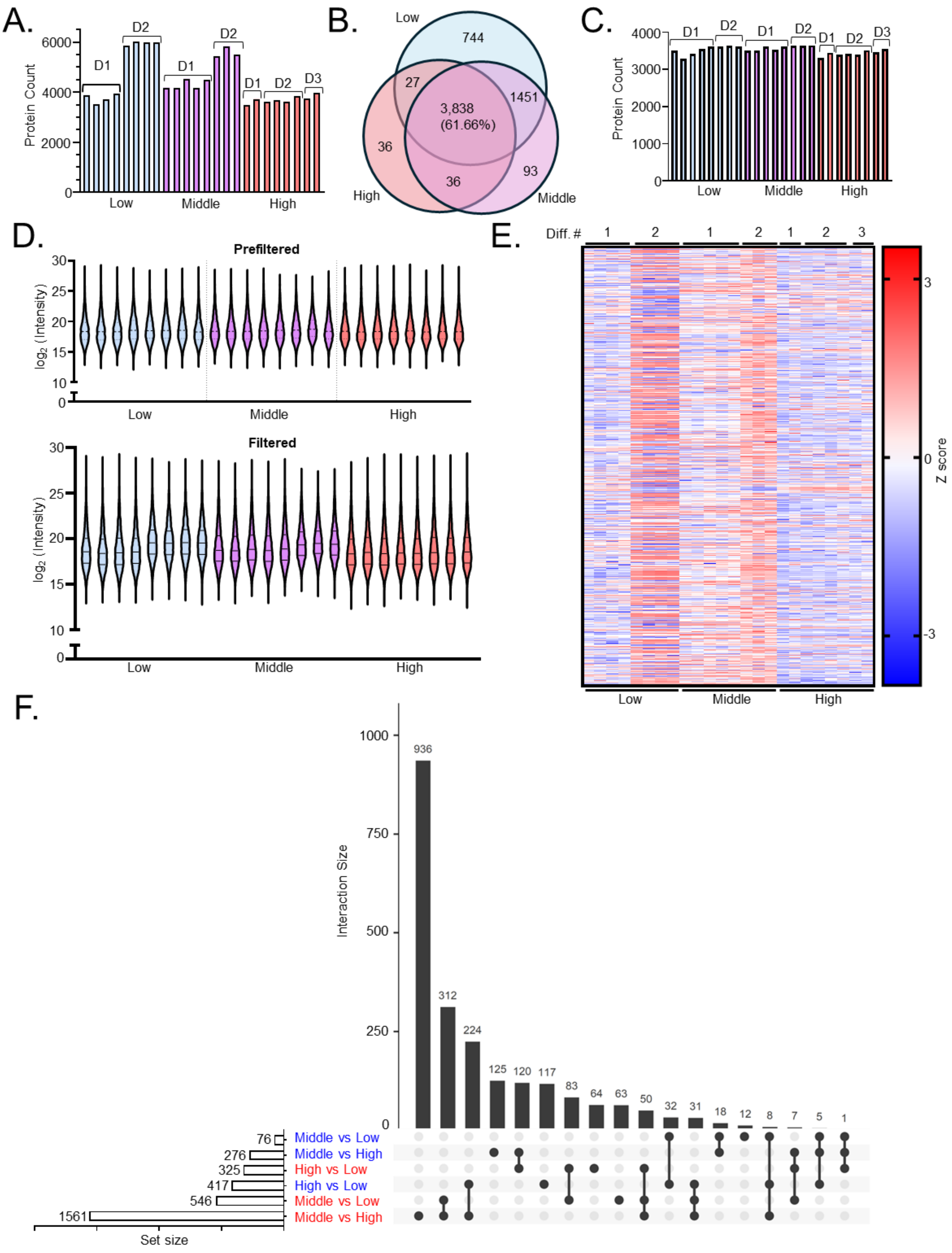
A.
Protein Count
6000
4000
2000
0
D1
D2
D1
D2
D1
D2
D3
Low
Middle
High
B.
Low
744
27
3,838
(61.66%)
1451
36
36
93
High
Middle
C.
Protein Count
4000
3000
2000
1000
0
D1
D2
D1
D2
D1
D2
D3
Low
Middle
High
D.
Prefiltered
log2 (Intensity)
30
25
20
15
10
0
Low
Middle
High
Filtered
log2 (Intensity)
30
25
20
15
10
0
Low
Middle
High
E.
Diff. #
1
2
1
2
1
2
3
3
0
-3
Z score
Low
Middle
High
F.
Interaction Size
1000
750
500
250
0
936
312
224
125
120
117
83
64
63
50
32
31
18
12
8
7
5
1
76
276
325
417
546
1561
Middle vs Low
Middle vs High
High vs Low
High vs Low
Middle vs Low
Middle vs High
Set size

**Supplemental Figure 1: Global proteomics of hiPSC-CMs with low-, medium- and high-QT-PGS. (A)** Number of proteins identified in 8 replicates for each cell line representing each QT-PGS. D1, D2, and D3 correspond to each differentiation respectively. **(B)** Venn diagram shows the overlap of identified proteins among all 3 cell lines. **(C)** Protein counts for each replicate after filtering for proteins identified in at least three samples and two separate differentiations. **(D)** Overall protein abundance ($log_2$ normalized) for each replicate before and after filtering as in C. **(E)** Z score heatmap of filtered data for global proteomics. Missing values were mean imputed based on each differentiation and cell line for visualization only. **(F)** Upset plot showing variable comparisons of the global proteomic data. Horizonal bars on the left show the total number of significant proteins in each individual comparison (set size). Each vertical bar represents the number of proteins significantly different between the indicated group comparisons (intersection size). Connected black dots below the bars show which group comparisons share those proteins. Red group labels indicate proteins increased in the first-named group, while blue labels indicate proteins decreased in the first-named group.

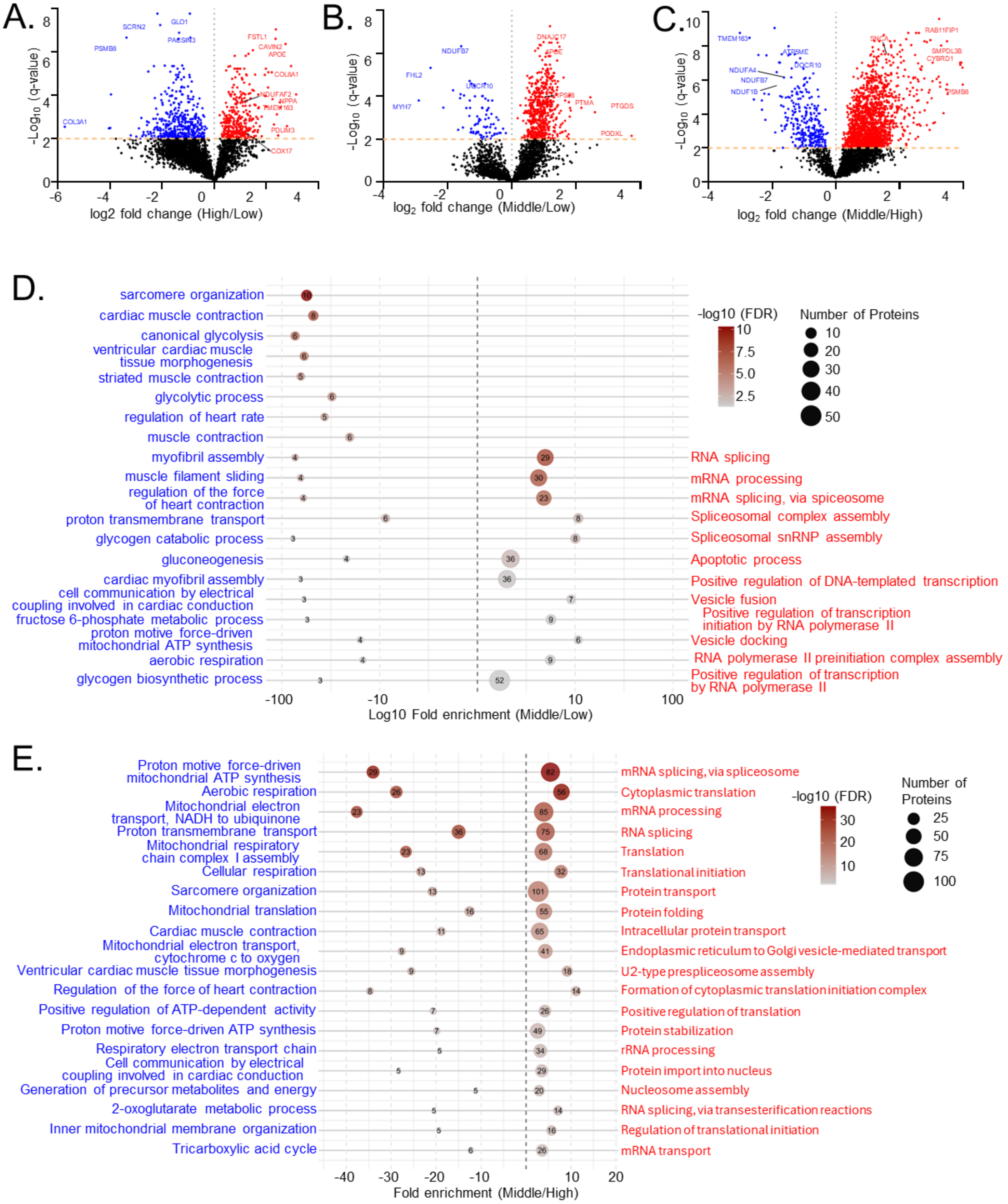


**Supplemental Figure 2. Mirrored bubble plots of significantly enriched GO biological processes identified from global proteomics analysis.**

**(A-C)** Volcano plots showing $\log_2$ fold change versus $-\log_{10}$ adjusted P values for three pairwise comparisons among low- medium- and high-QT-PGS groups. The orange dashed line indicates the FDR threshold for significance (FDR < 0.01; two-stage step-up procedure of Benjamini, Krieger, and Yekutieli).

Red = upregulated, blue = downregulated, black = not significant. Samples = 8 lysates per QT-PGS group from at least 2 separate differentiations (Low/Medium = 2 differentiations, High = 3 differentiations).
**(D)** and **(E)** show mirrored bubble plots representing enriched GO terms obtained using the DAVID functional annotation tool. GO enrichment was performed separately for proteins significantly lower (blue, left panels) or higher (red, right panels) for each QT-PGS comparison. The size of each bubble corresponds to the number of proteins associated with a given GO term, while the color intensity indicates the –log10(q-value) of enrichment significance, as shown on the color scale. Terms are ordered along the y-axis by their enrichment ranking.

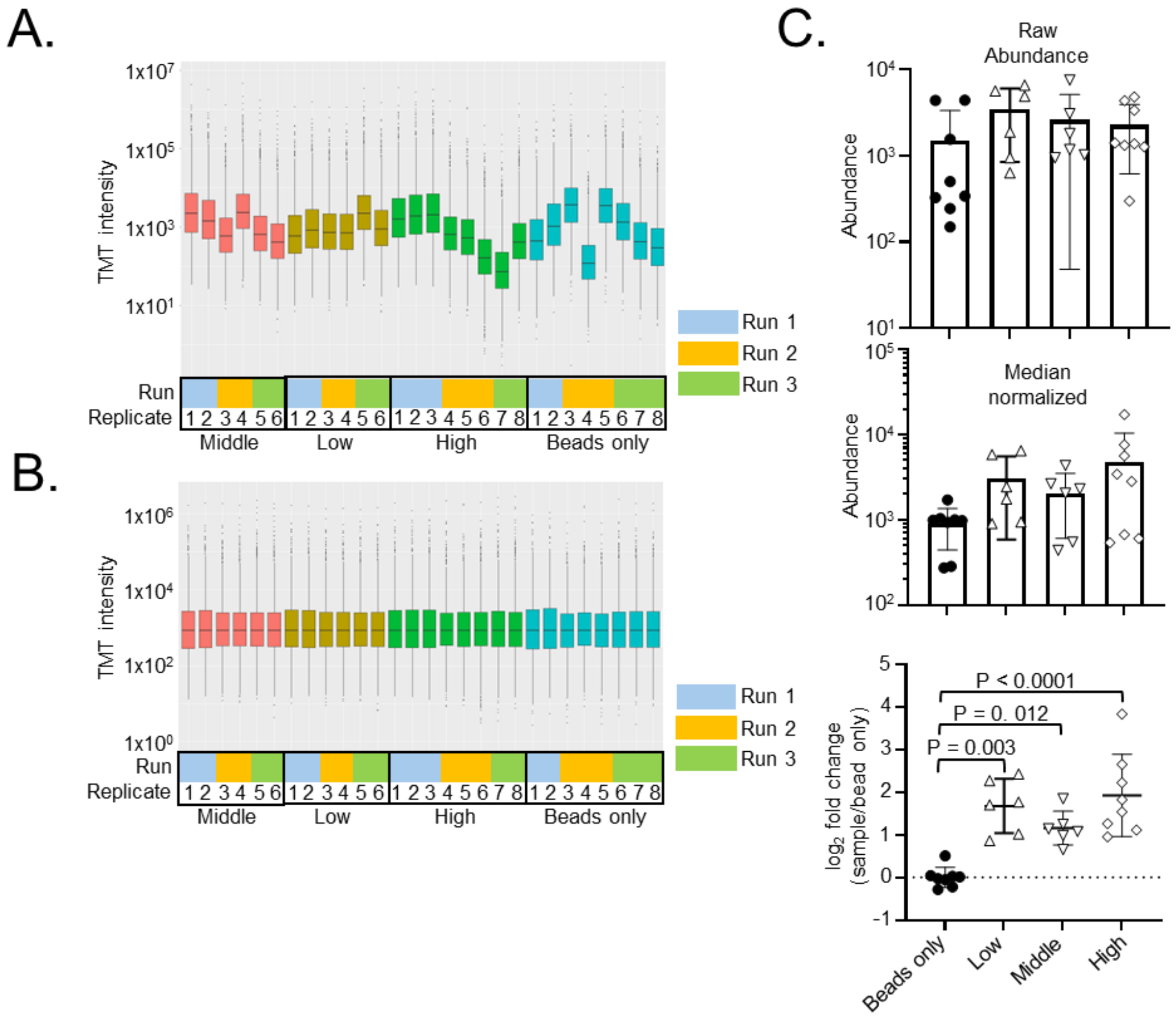


**Supplemental Figure 3: TMT and other AP-MS quantification overview. (A)** Box-and-whisker plots showing total TMT reporter ion intensities for all proteins across individual channels and runs. **(B)** Median normalization of protein abundance across all runs used for statistical analysis and quantification. **(C)** Kv11.1 (bait-protein) quantification showing raw abundance values, median-normalized data, and $log_2$-transformed values normalized to the bead-only-control in each run.

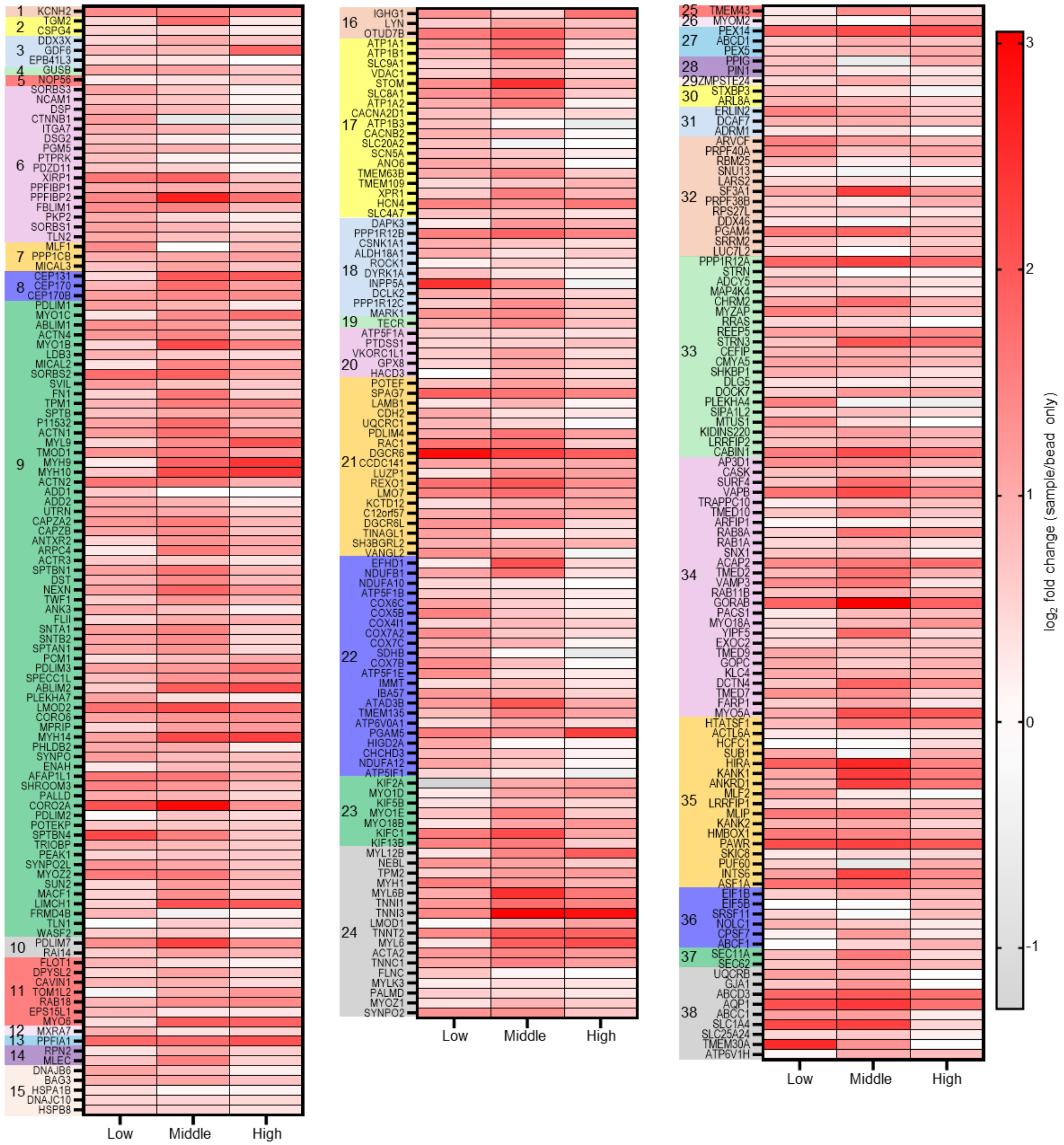


**Supplemental Figure 4: Heatmaps of Kv11.1 interactor abundance grouped by manual GO term assignment.** (1) Bait protein, (2) angiogenesis, (3) apoptosis, (4) Autophagy & Lysosomal Processing, (5) biogenesis, (6) cell adhesion, (7) cell cycle, (8) centrosome, (9) cytoskeleton, (10) differentiation, (11) endocytosis, (12) extracellular matrix, (13) focal adhesion, (14) GlycanProcessing, (15) Hsp40/70/90 & Other Chaperoning Activity, (16) immunity, (17) ion transport, (18) kinases and other enzymes, (19) lipid synthesis, (20) metabolism, (21) Miscellaneous, (22) mitochondrial function, (23) motor protein, (24) muscle contraction, (25) nuclear import/export, (26) oxidoreductase, (27) peroxisome, (28) Proline Isomerization, Hydroxylation, & Collagen Processing, (29) protein catabolism, (30) protein transport, (31) Protein Ubiquitination & Proteasomal Degradation, (32) RNA processing, (33) signaling, (34) trafficking, (35) transcription, (36) translation, (37) translocation, (38) transport.

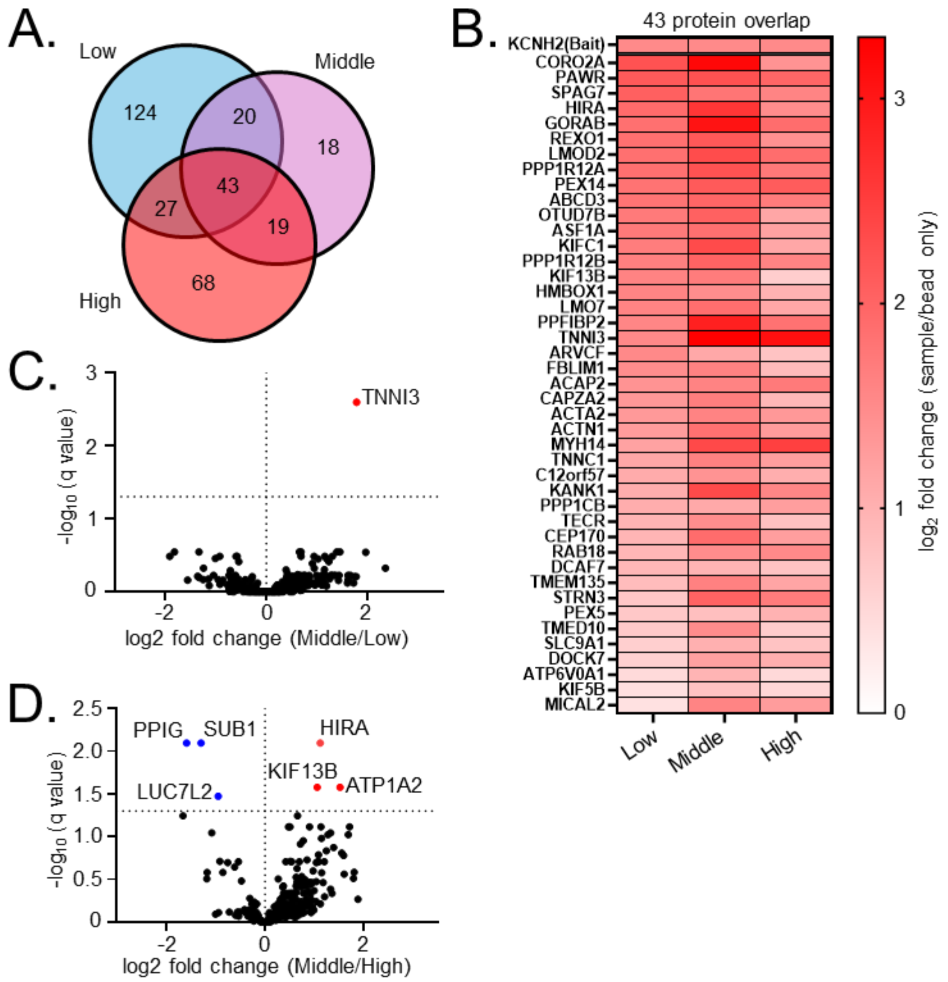


**Supplemental Figure 5: Kv11.1 core interactors and Middle QT-PGS comparisons. (A)** Venn diagram showing overlap of significant Kv11.1 interactors across QT-PGS groups. Out of 319 total enriched interactors, 43 interactors were significantly enriched in all three groups dictating a core Kv11.1 interactome. **(B)** Log2 fold change of 43 core interactors across QT-PGS groups. All abundances were normalized to KCNH2 bait control and compared to bead only pulldowns. **(C&D)** Volcano plots showing log2 fold change comparison of middle vs. low QT-PGS (C) and middle vs high (D) for all 319 Kv11.1 interactors.

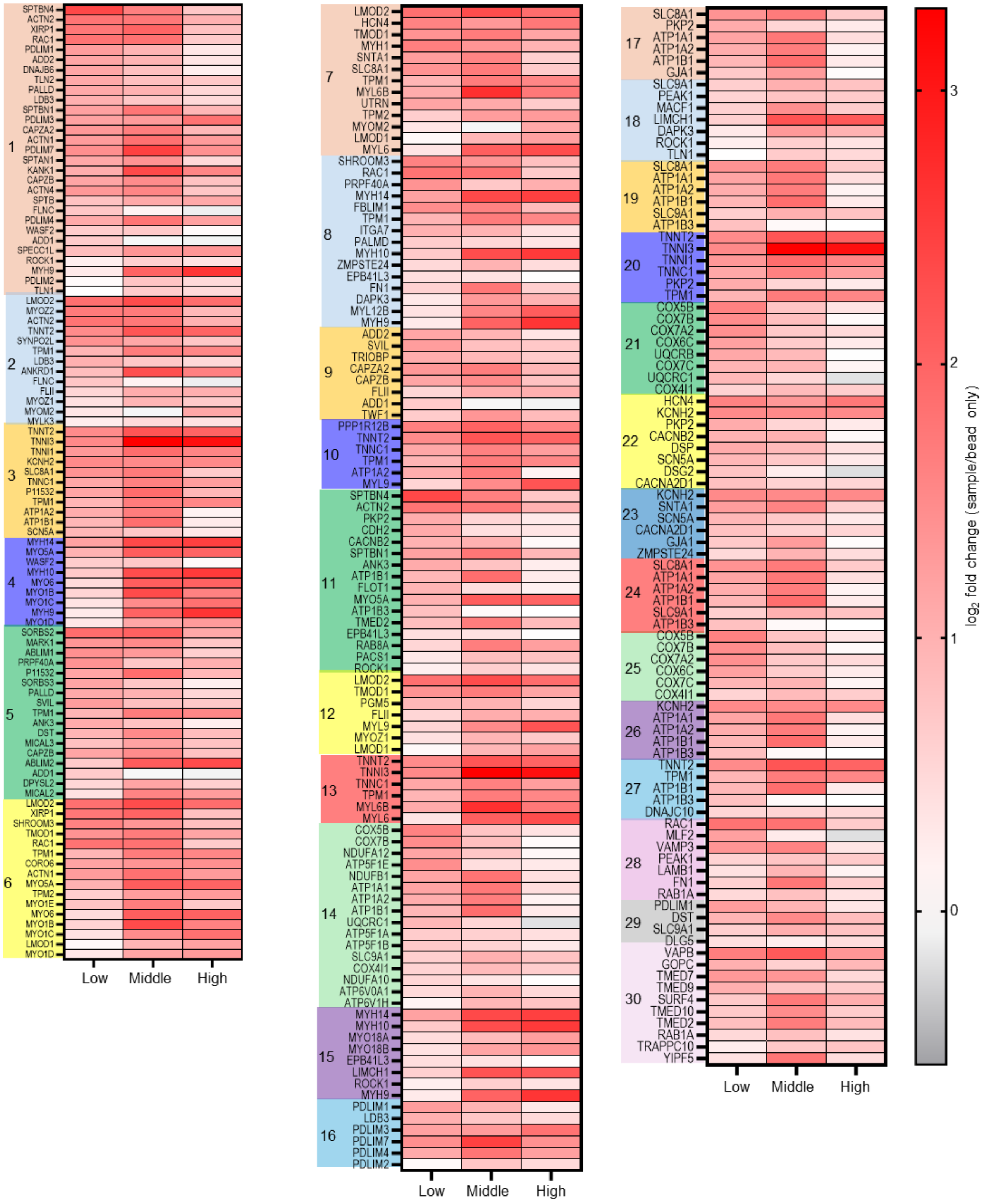

1
SPTBN4
ACTN2
XIRP1
RAC1
PDLIM1
ADD2
DNAJB6
TLN2
PALLD
LDB3
SPTBN1
PDLIM3
CAPZA2
ACTN1
PDLIM7
SPTAN1
KANK1
CAPZB
ACTN4
SPTB
FLNC
PDLIM4
WASF2
ADD1
SPECC1L
ROCK1
MYH9
PDLIM2
TLN1
2
LMOD2
MYOZ2
ACTN2
TNNT2
SYNPO2L
TPM1
LDB3
ANKRD1
FLNC
FLII
MYOZ1
MYOM2
MYLK3
3
TNNT2
TNNI3
TNNI1
KCNH2
SLC8A1
TNNC1
P11532
TPM1
ATP1A2
ATP1B1
SCN5A
4
MYH14
MYO5A
WASF2
MYH10
MYO6
MYO1B
MYO1C
MYH9
MYO1D
5
SORBS2
MARK1
ABLIM1
PRPF40A
P11532
SORBS3
PALLD
SVIL
TPM1
ANK3
DST
MICAL3
CAPZB
ABLIM2
ADD1
DPYSL2
MICAL2
6
LMOD2
XIRP1
SHROOM3
TMOD1
RAC1
TPM1
CORO6
ACTN1
MYO5A
TPM2
MYO1E
MYO6
MYO1B
MYO1C
LMOD1
MYO1D
Low
Middle
High
7
LMOD2
HCN4
TMOD1
MYH1
SNTA1
SLC8A1
TPM1
MYL6B
UTRN
TPM2
MYOM2
LMOD1
MYL6
8
SHROOM3
RAC1
PRPF40A
MYH14
FBLIM1
TPM1
ITGA7
PALMD
MYH10
ZMPSTE24
EPB41L3
FN1
DAPK3
MYL12B
MYH9
9
ADD2
SVIL
TRIOBP
CAPZA2
CAPZB
FLII
ADD1
TWF1
10
PPP1R12B
TNNT2
TNNC1
TPM1
ATP1A2
MYL9
11
SPTBN4
ACTN2
PKP2
CDH2
CACNB2
SPTBN1
ANK3
ATP1B1
FLOT1
MYO5A
ATP1B3
TMED2
EPB41L3
RAB8A
PACS1
ROCK1
12
LMOD2
TMOD1
PGM5
FLII
MYL9
MYOZ1
LMOD1
13
TNNT2
TNNI3
TNNC1
TPM1
MYL6B
MYL6
14
COX5B
COX7B
NDUFA12
ATP5F1E
NDUFB1
ATP1A1
ATP1A2
ATP1B1
UQCRC1
ATP5F1A
ATP5F1B
SLC9A1
COX4I1
NDUFA10
ATP6V0A1
ATP6V1H
15
MYH14
MYH10
MYO18A
MYO18B
EPB41L3
LIMCH1
ROCK1
MYH9
16
PDLIM1
LDB3
PDLIM3
PDLIM7
PDLIM4
PDLIM2
Low
Middle
High
17
SLC8A1
PKP2
ATP1A1
ATP1A2
ATP1B1
GJA1
18
SLC9A1
PEAK1
MACF1
LIMCH1
DAPK3
ROCK1
TLN1
19
SLC8A1
ATP1A1
ATP1A2
ATP1B1
SLC9A1
ATP1B3
20
TNNT2
TNNI3
TNNI1
TNNC1
PKP2
TPM1
21
COX5B
COX7B
COX7A2
COX6C
UQCRB
COX7C
UQCRC1
COX4I1
22
HCN4
KCNH2
PKP2
CACNB2
DSP
SCN5A
DSG2
CACNA2D1
23
KCNH2
SNTA1
SCN5A
CACNA2D1
GJA1
ZMPSTE24
24
SLC8A1
ATP1A1
ATP1A2
ATP1B1
SLC9A1
ATP1B3
25
COX5B
COX7B
COX7A2
COX6C
COX7C
COX4I1
26
KCNH2
ATP1A1
ATP1A2
ATP1B1
ATP1B3
27
TNNT2
TPM1
ATP1B1
ATP1B3
DNAJC10
28
RAC1
MLF2
VAMP3
PEAK1
LAMB1
FN1
RAB1A
29
PDLIM1
DST
SLC9A1
DLG5
30
VAPB
GOPC
TMED7
TMED9
SURF4
TMED10
TMED2
RAB1A
TRAPPC10
YIPF5
Low
Middle
High
3
2
1
0
log2 fold change (sample/bead only)

**Supplemental Figure 6: Heatmaps for Kv11.1 interactors grouped by top 30 DAVID assigned GO terms.** 1. Actin cytoskeleton organization, 2. Sarcomere organization, 3. Cardiac muscle contraction, 4. Actin filament based movement, 5. Cytoskeleton organization, 6. Actin filament organization, 7. Muscle contraction, 8. Regulation of cell shape, 9. Barbed end actin filament capping, 10. Regulation of muscle contraction, 11. Protein localization to plasma membrane, 12. Myofibril assembly, 13. Muscle filament sliding, 14. Proton transmembrane transport, 15. Actomyosin structure organization, 16. Muscle structure development, 17. Cell communication by electrical coupling involved in cardiac conduction, 18. Regulation of focal adhesion assembly, 19. Sodium ion export across plasma membrane, 20. Ventricular cardiac muscle tissue morphogenesis, 21. Cellular respiration, 22. Regulation of heart rate by cardiac conduction, 23. Regulation of ventricular cardiac muscle cell membrane repolarization, 24. Intracellular sodium ion homeostasis, 25. Mitochondrial electron transport cytochrome c to oxygen, 26. Membrane repolarization, 27. Positive regulation of ATP dependent activity, 28. Substrate adhesion dependent cell spreading, 29. Maintenance of cell polarity, 30. Endoplasmic reticulum to Golgi vesicle mediated transport.

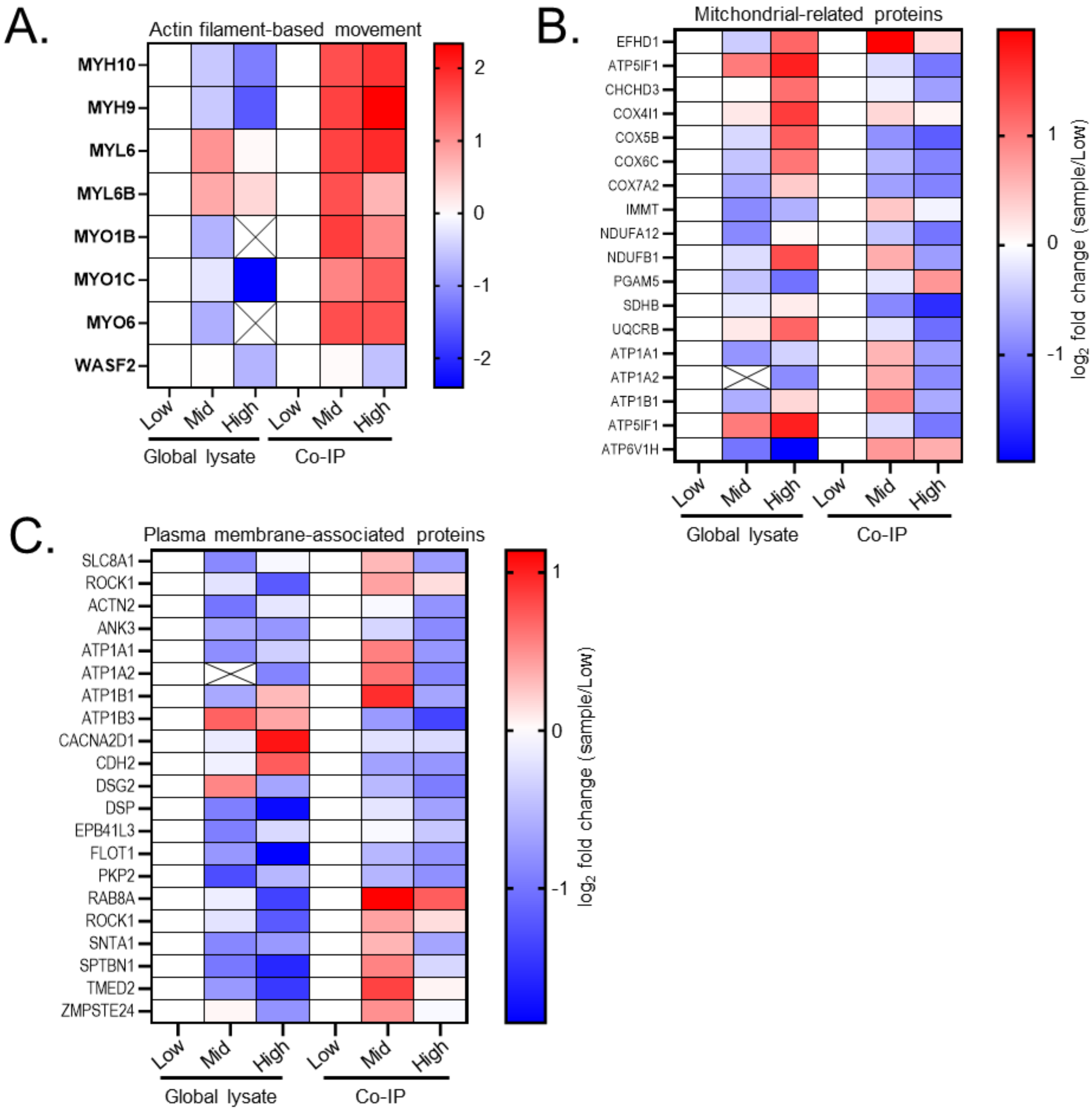


**Supplemental Figure 7: Heatmaps showing comparison of global proteomics to Kv11.1 enriched interactors. (A-C)** Heatmaps of protein abundances for (A) actin filament-based movement, (B) mitochondrial-related proteins and (C) Plasma membrane associated proteins from global lysates measured by data-independent acquisition mass spectrometry and from affinity enriched Kv11.1 interactors (Co-IP) quantified with data-dependent acquisition with TMT labeling. X marks proteins not identified in at least 3 samples in the global proteomics dataset. Protein abundance was normalized to the low QT-PGS group for each acquisition method and shown as log2 fold-change.
